# A versatile multimodal optical modality based on Brillouin light scattering and photoacoustic effect

**DOI:** 10.1101/2023.03.10.532144

**Authors:** Chenjun Shi, Yan Yan, Mohammad Mehrmohammadi, Jitao Zhang

## Abstract

Multimodal optical imaging techniques are useful for various applications, including imaging biological samples for providing comprehensive material properties. In this work, we developed a new modality that can measure a set of mechanical, optical, and acoustical properties of a sample at microscopic resolution, which is based on the integration of Brillouin (Br) and photoacoustic (PA) microscopy. The proposed multimodal imaging technique not only can acquire co-registered Br and PA signals but also allows us to utilize the sound speed measured by PA to quantify the sample’s refractive index, which is a fundamental property of the material and cannot be measured by either technique individually. We demonstrated the colocalization of Br and time-resolved PA signals in a synthetic phantom made of kerosene and CuSO_4_ aqueous solution. In addition, we measured the refractive index of saline solutions and validated the result against published data with a relative error of 0.3 %. This multimodal Br-PA modality could open a new way for characterizing biological samples in physiological and pathological conditions.

---

Photoacoustic (PA) imaging is a rapidly growing modality in biomedical research due to its ability to acquire high-contrast functional and molecular images of biological samples at relatively large depths compared to other optical imaging techniques [1-3]. In PA imaging, the absorption of light energy (i.e., short laser pulses) by chromophores (i.e., endogenous or exogenous) generates acoustic waves caused by rapid thermoelastic expansion [1]. The consequent acoustic waves can be collected by acoustic detectors such as an ultrasound transducer or a hydrophone to form PA images. The amplitude of the detected PA signal represents the optical absorption and scattering of the object. In addition, in PA microscopy, the speed of sound (SOS) can be obtained by calculating the time-of-flight (TOF) of the PA signal with prior knowledge of the physical distance between the detector and the location of the excitation beam.

Confocal Brillouin microscopy is an emerging optical modality for quantifying the mechanical properties of materials with a diffraction-limit resolution [4-7]. The principle of Brillouin microscopy is based on spontaneous Brillouin scattering, where the interaction of the incident light and the thermal acoustic phonons within the sample introduces a frequency shift (i.e., Brillouin shift *ω*_*B*_) to the scattered light. The Brillouin shift at 90° geometry is physically determined by 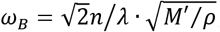, where *λ* is the laser wavelength, *n* is the refractive index, *ρ* is the sample’s density, and *M*′ is the elastic longitudinal modulus. With a known refractive index and density, the sample’s mechanical properties can be directly quantified with the Brillouin shift measured by a Brillouin spectrometer [8].

In the past two decades, the Brillouin microscope has been demonstrated for non-invasive mechanical imaging of cells and tissues, indicating its promising applications in ocular diseases, cancer metastasis, cellular biomechanics, and developmental biology [9-14]. Currently, Brillouin microscope mostly uses the Brillouin shift to estimate the relative change of longitudinal modulus, with the assumption that the ratio of refractive index and density 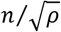 is approximately constant in many physiological processes [5, 11, 15-18]. However, for direct and accurate quantification of longitudinal modulus, measurement of the refractive index and density is required. Furthermore, since the density of many biological materials can be calculated from the refractive index based on a two-substance mixture model [19-21], colocalized measurement of refractive index and Brillouin shift will allow the direct quantification of longitudinal modulus. Very recently, dual-geometry Brillouin microscopy [22] and a multimodal modality that combines Brillouin microscopy with optical diffraction tomography [23] have been demonstrated for this purpose.

In this work, we report for the first time a multimodal optical modality by combining Brillouin and PA microscopy. In this hybrid modality, two laser beams were coupled into a common optical path for collecting the Brillouin and time-resolved PA signals from the same spot simultaneously. Therefore, this new modality allows us to acquire the colocalized optical, acoustic, and mechanical properties of the material for comprehensive characterization. Intriguingly, using the SOS (*V*_*S*_) measured by the PA signal, we can directly quantify the refractive index from the measured Brillouin shift based on the relationship of 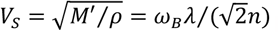, which further allows us to derive the longitudinal modulus based on the two-substance mixture model. It is worth highlighting that, this additional information can only be obtained by the integrated system rather than by Brillouin or PA modality individually.

Figure 1 shows the schematic of the multimodal Brillouin-PA microscopic imaging system. A 780-nm continuous-wave (CW) laser (DL pro, Toptica) was used as the light source for Brillouin scattering. The laser beam first went through an isolator to reject possible back-reflection. A variable ND filter was used to adjust the output power of the laser, and a half-wave plate (HWP1) was used to adjust the polarization state of the beam to be p-polarized. Using a pair of lenses (L1, *f* = 30 mm and L2, *f* = 150 mm), the diameter of the laser beam was expanded from 2.38 mm to 11.90 mm. After passing through the second polarized beam splitter (PBS2), the mirror M1, and a half-wave plate (HWP3), the laser beam was focused into the sample cell (1-cm square cuvette) by lens L3 (*f* = 75 mm), with a focused spot of 6.26 μm. An in-house-built Brillouin spectrometer was used to collect the Brillouin signal at 90° geometry. The light source for PA excitation, emitted from a tunable nanosecond pulsed laser (OPOTEK, Phocus Mobile), was tuned to the wavelength of 780 nm and had a beam diameter of ∼8 mm. Since the emitted light from the pulsed laser is randomly polarized, a polarized beam splitter (PBS1) was first used to obtain a linearly p-polarized beam. After adjusting the polarization orientation of the PAbeamto s-polarized with ahalf-wave plate (HWP2), the PA beam was coupled into the optical path of the Brillouin beam using PBS2 and focused into the sample cell with a spot size of 9.31 μm in diameter. The half-wave plate (HWP3) was used to further adjust the Brillouin beam and PA beam to s-polarized and p-polarized, respectively. Since 90° scattering geometry is sensitive to s-polarized light but has no response to p-polarized light [24], our design can generate a strong Brillouin signal while avoiding any crosstalk from the PA excitation beam.

**Fig. 1.**
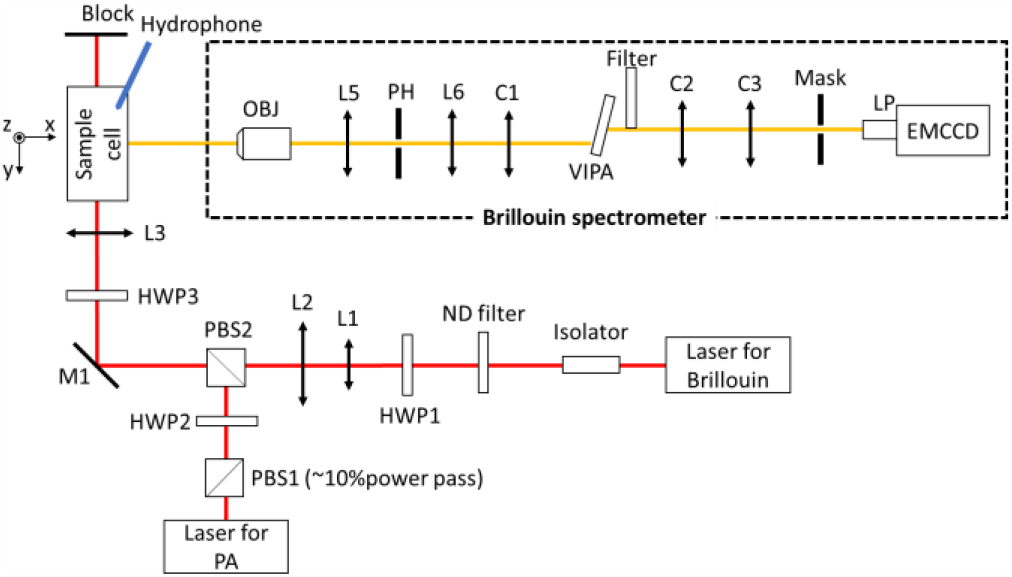
Schematic of the multimodal modality based on Brillouin and PA imaging. HWP1-HWP3: half-wave plates; PBS1-PBS2: polarized beam splitter; L1-L5: lenses; OBJ: objective; PH: pin hole; C1-C3: cylindrical lenses; LP: lens pair.

The generated PA signal was detected by a needle hydrophone (Onda, HNP-0400, 1 - 20 MHz) that was inserted into the sample cell and closely distanced from the focal point (∼ 4 mm). Meanwhile, the Brillouin signal was measured by a virtually imaged phased array (VIPA)-based Brillouin spectrometer. Upon being collected by an objective (OBJ, 4x/0.1), the Brillouin signal passed through a confocal unit consisting of two lenses and an adjustable pinhole (L5, *f* = 19 mm; L6, *f =* 80; and PH: tunable pinhole), through which the Brillouin signal out of the focal plane was rejected. A cylindrical lens (C1, *f* = 200 mm) was used to couple the Brillouin beam into the VIPA etalon (FSR=15 GHz, LightMachinery). A Filter (continuous ND filter, Thorlabs) was used for apodization. The output beam of the VIPA was reshaped by the cylindrical lenses C2 (*f* = 150 mm) and C3 (*f* = 75 mm) and then projected onto the mask. A lens pair (LP, 1:1, f = 30.0 mm) was usedtoimage the Brillouin spectrum onto an electron-multiplying charge-coupled devices (EMCCD) camera (iXon, Andor). Before experiments, the spectrometer was calibrated using standard materials (i.e., water and methanol) based on established protocol [7].

To evaluate the colocalization of the multimodal PA and Brillouin imaging system, we conducted an experiment in which a synthetic phantom (Fig. 2(a)) made of 1% copper sulfate solution (CuSO_4_) and kerosene was scanned to acquire both PA and Brillouin profiles across the sample. For the Brillouin measurement, the laser power was 27 mW at the focal plane, and the acquisition time of the spectrometer was 50 ms. For PA measurement, the average energy of the laser pulse was 1.4 mJ at a repetition rate of 10 Hz. Fig. 2 (b) and (c) show the representative Brillouin shift and PA amplitude for CuSO_4_ solution and kerosene, respectively. The sample cell was carried by a translation stage and was scanned manually along the z-direction with a step size of 0.25 mm and a total travel range of 1.75 mm. At each position, 100 frames of the Brillouin and the PA signals were collected for calculating the average Brillouin shift and PA peak-to-peak amplitude. Fig. 3 shows the colocalized Brillouin and PA signals across the interface of two materials. The co-registered contrast trends for Brillouin and PA measurements confirmed that the Brillouin beam and the PA beam share the same focal point in the sample. At the transition zone, the increase of PA signal amplitude and the decrease of Brillouin shift is mainly due to the beam distortion caused by the curved interface of two liquids (Fig.2(a)). The step size is limited by the resolution of the manual translational stage.

**Fig. 2.**
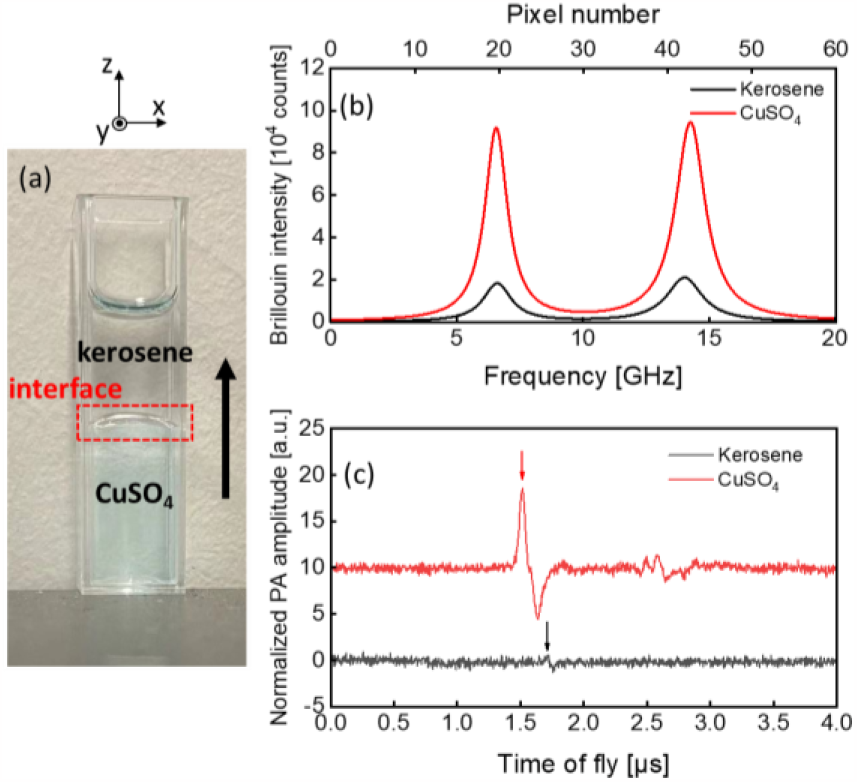
Experimental sample and signals. (a) Photo of the stratified sample with the scan direction. (b) Raw Brillouin and (c) PA signal (single measurement) of CuSO_4_ and kerosene. Arrows indicate the PA signal peaks.

**Fig. 3.**
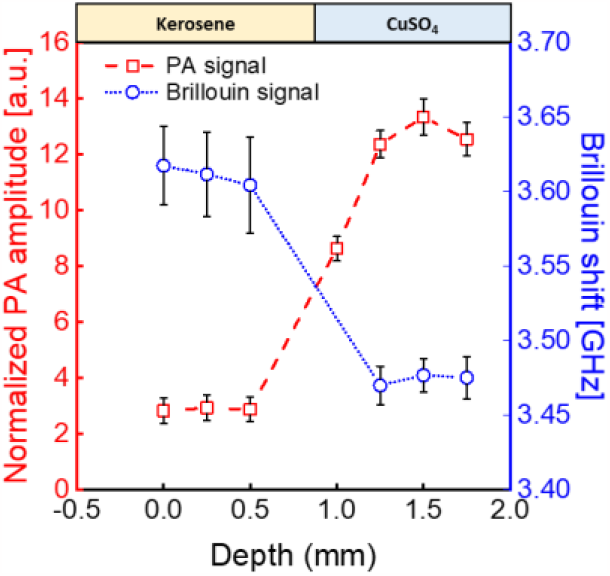
Result of 1D scanning. Vertical 1-D profiling of the stratified CuSO_4_-kerosene sample based on (a) Brillouin shift and (b) PA amplitude. Error bars are standard deviation for 100 times measurement.

We further explored the utility of the developed multimodal modality by directly measuring the refractive index. To do so, we prepared three saline solutions: the weight/weight concentrations are 0% (deionized water), 4.76%, and 16.67%, respectively. The SOS (*V*_*S*_) was measured through consecutive PA acquisitions, where the distance between the detecting hydrophone and the incident beam in the sample was changed with the pre-set value of Δ*d*. The time delay between the TOFs of these two consecutive PA acquisitions was measured as Δ*t* (Fig. 4(a)), allowing the calculation of sound speed by *V*_*S*_ = Δ*d*/Δ*t*. For each sample, the transducer was moved to four positions in sequence, with a distance of 0.25 mm in each movement (moving accuracy: ±0.02 mm). At each position, the PA signal was averaged by 100 times, and the primary positive peak was used to quantify the arrival time. The value of *V*_*S*_ was then calculated from all datasets by linear regression using the least squares method (Fig. 4 (b)). Simultaneously, Brillouin shift *ω*_*B*_ of the sample was measured as described earlier. Therefore, the refractive index of the sample can be calculated by

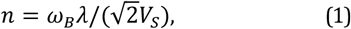

with *λ* = 780 nm. For each sample, we repeated the measurements three times for both PA and Brillouin, and the averaged values of the refractive index with the standard error of the mean were shown in Fig. 5. The measured values were compared with the results from Fiore *et al*. using dual-geometry Brillouin spectroscopy at 532 nm [22] and from Esteban *et al*. using a fiber-optic sensor at 780 nm [25]. Taking the average of the literature data (Fiore’s data were corrected based on water’s refractive index at 780 nm and 20.7°C) as a reference, our results show a discrepancy of 0.0041 (0.31%), 0.0029 (0.22%), and 0.0042 (0.31%), respectively, suggesting a good agreement.

**Fig. 4.**
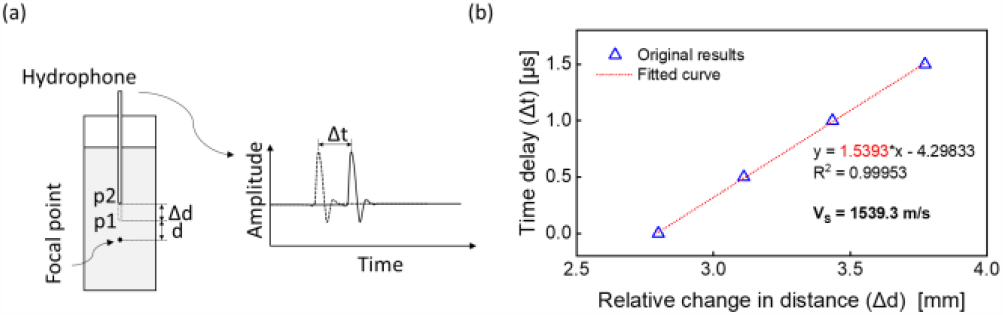
Speed of sound measurement. (a) the schematic for measuring SOS; (b) the regression results from the measurement on 4.76% saline.

**Fig. 5.**
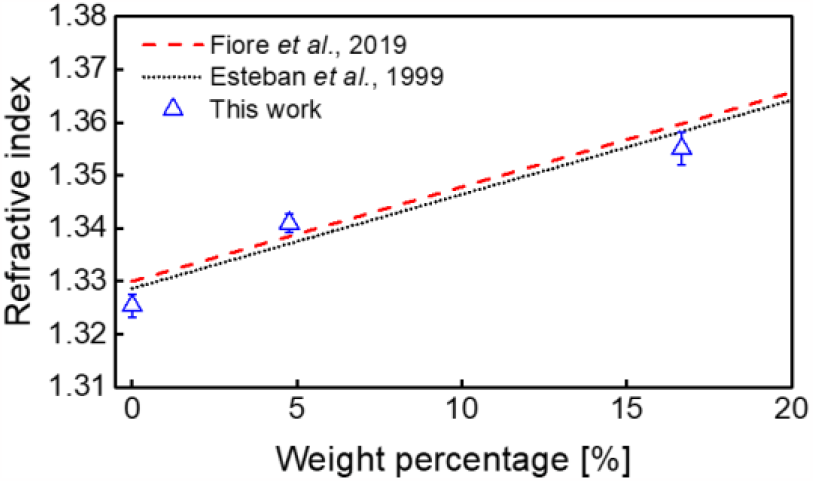
Comparison between measured refractive index and the literature. Error bar represents standard error of the mean.

Since Brillouin microscopy probes the acoustic phonon at the GHz frequency band while the PA signal is in the MHz band, the possible acoustic dispersion of the sample may introduce an artifact to the measured refractive index. To estimate the acoustic dispersion, we calculated the phonon velocity at GHz derived from the Brillouin measurements using the reported value of the refractive index [25] and compared it with PA measurements at MHz. The result is summarized in Table 1, where the relative discrepancy between the two datasets is within 0.4%, indicating that the acoustic dispersion in our samples is negligible. This is also consistent with several previous studies in which the SOS in pure water and saline has a variety of less than 0.3% as the frequency spans from 0.5 MHz to 1.5 GHz [26-30]. In addition, a linear behavior (i.e., no dispersion) of the sound wave has been found in many (bio)polymers [22], indicating the potential application of our multimodal modality in biomaterials and biological samples. For materials in which acoustic dispersion is prominent, prior knowledge of dispersion behavior is needed for accurate measurement of the refractive index.

**Table 1.**
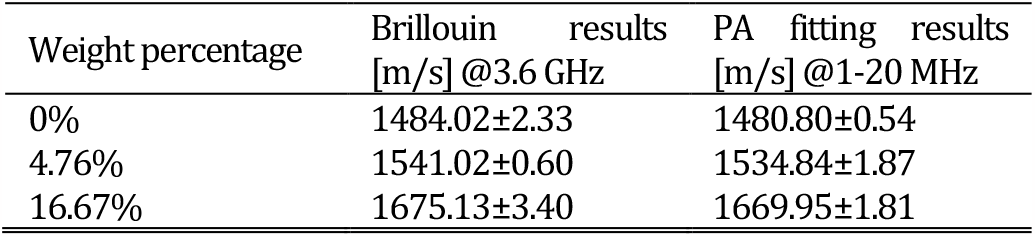
Comparison of SOS results calculated by Brillouin and PA (with standard errors).

Here we used a homogenous sample to validate the feasibility of refractive index measurement. In fact, our multi-modal imaging technique can also be used for inhomogeneous material such as multi-layered samples. In this case, the thickness of each layer can be first obtained from Brillouin images. While keeping the PA detector still, the movement of the beam spot into different layers will introduce time delay to the PA signal, which is a function of SOS for each layer. Therefore, the SOS of each layer can be derived by solving a set of multivariate equations. Together with the acquired Brillouin image, the refractive index of each layer can be measured. While the current prototype utilized two separate laser sources for generating Brillouin and PA signals, a single laser source could be used for both BA and Br microscopy in case a pulse laser with narrow linewidth and nanosecond pulse width is available. In the present work, the optical setup was demonstrated in transmission mode. It can be easily modified into a reflection mode byintegrating with the existing designs of inverted confocal Brillouin microscopy and PA microscopy[31], allowing better accessibility for biomedical applications.

In summary, we proposed a versatile multimodal optical imaging modality based on the combination of Brillouin microscopy and PA imaging. The integrated system with a common focal point for both Brillouin and PA beam was designed to simultaneously obtain the mechanical, optical, and acoustical properties of materials. Importantly, the integrated system can directly quantify the sample’s refractive index, which is not accessible by each individual technique. Together, the proposed versatile multimodal Brillouin and PA imaging system has the potential to provide quantitative measurements of multiple parameters for biomedical research.

## Funding

National Institutes of Health (K25HD097288); National Institutes of Health (R01EB030058), University Research Grant of Wayne State University; Richard Barber Interdisciplinary Research Program.

## Acknowledgments

The authors would like to thank Dr. Matthew O’Donnell from the University of Washington for discussions and valuable feedback on sound speed measurement studies presented in this work. We also want to thank Dr. Antonio Fiore from HHMI Janelia Research Campus for providing raw data of refractive index measurements with dual-geometry Brillouin microscopy.

## Disclosures

A provisional patent application related to this research has been filed by Wayne State University patent office for J.Z., C.S., Y.Y., and M. M..

## Data availability

Data underlying the results presented in this paper can be obtained from the authors upon reasonable request.

